# Icecream: High-Fidelity Equivariant Cryo-Electron Tomography

**DOI:** 10.1101/2025.10.17.682746

**Authors:** Vinith Kishore, Valentin Debarnot, Ricardo D. Righetto, Benjamin D. Engel, Ivan Dokmanić

## Abstract

Cryo-electron tomography (cryo-ET) visualizes 3D cellular architecture in near-native states. Recent deep-learning methods (CryoCARE, IsoNet, DeepDeWedge, CryoLithe) improve denoising and artifact correction, but performance remains limited by very low signal-to-noise ratio, a restricted angular range (‘missing wedge’), and the lack of ground truth. Here, we present Icecream, which follows the broad template of earlier self-supervised approaches, but treats symmetry in a way consistent with the recent equivariant imaging framework (Chen et al., 2021). Coupled with several engineering refinements, including mixed-precision arithmetic, Icecream achieves substantially better denoising and more reliable missing-wedge recovery, while reducing training and inference time relative to comparable baselines. Across diverse experimental datasets, we observe consistent gains in reconstruction quality, both visually and as quantified by Fourier shell correlation (FSC). Our framework extends to any tomography problem that provides two statistically independent reconstructions of the same volume; in cryo-ET these are obtained by dose splitting or angular partitioning of the tilt series.

## 1 Introduction

Cryo-electron tomography (cryo-ET) has become a central technique in structural biology, enabling direct visualization of macromolecular assemblies in their native cellular environment. Its ability to provide three-dimensional structural information at nanometer resolution without the need for crystallization has made it an essential tool for studying complex and heterogeneous biological systems such as organelles, pathogens, and large protein complexes. Cryo-ET complements single-particle cryo-EM by offering spatial context and capturing the structural diversity of biological specimens in situ (Navarro, 2022; McCafferty *et al*., 2024).

Despite its rapidly growing importance, cryo-ET remains extremely challenging from a computational and signal processing perspective. It involves imaging tiny, delicate, radiation-sensitive samples using high-energy electrons while tilting the sample with a mechanical stage. To minimize beam-induced damage, only a limited number of projections can be collected, each with a low dose. This results in a very low signal-to-noise ratio (SNR), with complex noise statistics that are difficult to model or filter without distorting meaningful signal (Vulović *et al*., 2013).

Further problems arise from mechanical limitations and sample geometry which prevent imaging across the full ±90^◦^ tilt range, leaving a significant portion of the Fourier space unconstrained. This so-called *missing wedge* results in anisotropic artifacts, most notably streaking and smearing, that degrade the resolution of reconstructed volumes (Shkolnisky & Singer, 2012; Chen *et al*., 2016).

A variety of post-processing and reconstruction techniques have been proposed to address these challenges. Classical approaches include filtering (Feldkamp *et al*., 1984; van der Heide *et al*., 2007; Frangakis, 2021), sub-tomogram averaging (Tegunov *et al*., 2021; Burt *et al*., 2024), and model-based regularization (Gilbert, 1972; Andersen & Kak, 1984; Deng *et al*., 2016; Yan *et al*., 2019). More recently, deep learning has emerged as a powerful tool for improving reconstruction quality, particularly in denoising and missing-wedge artifact removal.

### 1.1 Self-supervised learning

Our proposed method—Icecream—is self-supervised, meaning that it does not require clean ground-truth volumes to train. Self-supervised approaches are among the strongest successful applications of deep learning in cryo-ET reconstruction, where ground truth data does not exist. Perhaps the most common is CryoCARE (Buchholz *et al*., 2019) which builds on the more general Noise2Noise principle (Lehtinen *et al*., 2018): leveraging independent noisy observations to train a denoising neural network without clean supervision.

Self-supervised methods have also been developed for missing wedge correction. IsoNet exploits the fact that the orientation of the missing wedge in Fourier space doesn’t depend on the orientation of the volume (Liu *et al*., 2022). By training on rotated versions of the input sub-tomograms, IsoNet can partially recover the missing spectral regions using the observed wedge across different orientations. While it also includes a denoising mechanism, practitioners often apply CryoCARE before passing to IsoNet as this significantly improves denoising. To address the separate denoising and missing wedge completion, DeepDeWedge takes a more principled approach and combines Noise2Noise with the rotation-aware strategy used in IsoNet (Wiedemann & Heckel, 2024). This leads to a unified framework, but it comes at the cost of significantly increased training time. A detailed comparison of our approach with DeepDeWedge and CryoCARE+IsoNet is provided in Appendix A.

### 1.2 Supervised learning

A drawback of self-supervised methods is that they need to be trained from scratch for each new acquisition. Despite several attempts to train such models on large-scale datasets, their performance degrades when applied to unseen samples or different acquisition conditions (Wiedemann & Heckel, 2024). To overcome this challenge, we recently introduced CryoLithe (Kishore *et al*., 2025), the first supervised deep neural network designed to generalize across acquisitions, which eliminates the need for retraining. CryoLithe is robust to unseen observations thanks to localized learning strategies (Khorashadizadeh *et al*., 2025*a*; Khorashadizadeh *et al*., 2025*b*), but it still implicitly relies on self-supervised learning to generate high-quality training targets. Hence improving the quality of self-supervised reconstruction methods remains crucial.

### 1.3 Group equivariance

Chen et al. (Chen *et al*., 2021; Chen *et al*., 2022) introduced a theoretical framework for solving inverse problems by leveraging group equivariance as a form of regularization. When the group is the rotation group, equivariance is similar to the rotation property leveraged by IsoNet and DeepDeWedge. Indeed, most successful self-supervised methods in cryo-electron microscopy and tomography rely on augmenting the training dataset by rotated versions of the volume to process and a ‘manual’ masking of the missing wedge (Liu *et al*., 2022; Wiedemann & Heckel, 2024; Liu *et al*., 2025). Training on rotation-augmented data encourages the composition of the network with the forward operator to be rotation-equivariant, but the way this is achieved in prior work on cryo-ET is different from the equivariant imaging framework of Chen et al. We note that it is also possible to design neural networks that are equivariant to given groups by design (Chaman & Dokmanic, 2021; Chaman & Dokmanić, 2021; Herbreteau *et al*., 2023).

### 1.4 Contributions and outline

In this paper, we connect equivariant imaging as introduced by Chen et al. (Chen *et al*., 2021; Chen *et al*., 2022) with cryo-ET. This yields Icecream which achieves state-of-the-art denoising and missing wedge correction. We demonstrate Icecream on a variety of real experimental datasets with very diverse spatial statistics. We also optimize it so that it is faster and more memory efficient than existing self-supervised methods, while yielding better reconstructions.

The paper is organized as follows. Section 2 describes the acquisition model, the proposed method, and its training procedure. Section 3 presents quantitative and visual results on real data and demonstrates the interest of Icecream to accelerate the reconstruction process. Finally, Section 4 discusses the implications and limitations of the approach. Additional experiments and details about baseline methods are provided in the Appendix.

Icecream is openly available under the MIT License. The source code and the documentation can be found at github.com/swing-research/icecream.

## 2 Methods

We propose to learn a function *f*_*ϕ*_ — here a deep neural network — parameterized by *ϕ* ∈ Φ ⊂ ℝ^*P*^, which maps noisy and missing-wedge-degraded sub-tomograms to volumes free of artifacts. To achieve this, we combine two consistency criteria that jointly constrain the network to perform effective denoising and missing wedge correction: Noise2Noise and equivariance.

### 2.1 Acquisition model

Cryo-ET reconstruction algorithms estimate a three-dimensional (3D) volume density from a series of aligned two-dimensional (2D) projections, known as a tilt-series. We work with sub-tomograms and let 𝒳 be the set of sub-tomograms of size *N* × *N* × *N* we aim to recover, extracted from a larger clean (unavaialable) tomogram. We aim at improving the degraded measurements in two ways: (i) by estimating the missing wedge coefficients using explicit geometric priors, and (ii) by denoising the observed (visible wedge) coefficients by exploiting the statistical independence of different noisy observations of the underlying signal.

We introduce the spectral masking operator *A* which zeroes out the Fourier coefficients corresponding to the missing wedge. Denoting the limited-view tomographic forward operator by *F*, we can write *A* = *F* ^+^*F*, where (·)^+^ is the Moore–Penrose pseudoinverse (so *A* is an orthogonal projection on the observed wedge). In the following, we will apply *A* to sub-tomograms rather than to the entire volume. This is only an approximation of the true missing wedge operator, since cropping a sub-tomogram is equivalent to convolving its Fourier coefficients with a sinc function. We nonetheless adopt this approximation as it is computationally far more efficient.

Noise in cryo-ET is complex and arises from a combination of sources, including detector characteristics, sample-induced scattering, background signal, and electronic noise. Due to this complexity, a precise parametric model of the noise is intractable (Quinto *et al*., 2009; Yang *et al*., 2024) and we simply model it by a random operator P : ℝ^*N×N×N*^ → ℝ^*N×N×N*^, where the sub-tomograms are perturbed independently according to a fixed (perhaps unknown) distribution.

Modern direct detectors record multiple frames for every tilt angle (Faruqi & Henderson, 2007). These frames are then usually averaged after motion compensation in order to increase the signal-to-noise ratio. Recent deep learning approaches such as Cryo-CARE leverage the independence of noise realizations across the frames to outperform simple averaging. The effectiveness of these approaches relies on a trade-off between the amount of signal in the observations and the number of realizations. Similarly to Cryo-CARE, we also divide observations into two independent sets, either by splitting the dose as enabled by direct detectors, or by splitting the tilt-series along the tilt angles. We model this by assuming that for each volume *x* ∈ 𝒳, we get two independent observations

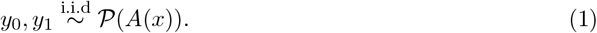

We let

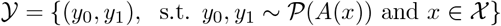

denote the set of observations corresponding to 𝒳.

In our implementation, we extract the sub-tomograms *y* ∈ 𝒴 following IsoNet’s procedure (Liu *et al*., 2022), after a tomogram-level normalization. This involves extracting the parts of the tomograms that are likely to contain structures of interest, and avoiding the parts that contain only ice.

### 2.2 Denoising and data consistency

We first ensure that the reconstruction is consistent with the measurements. For that, we follow the Noise2Noise principle which is among the most effective denoising techniques in cryo-ET. It amounts to minimizing the following cost function:

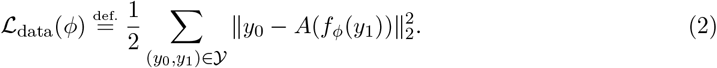

Notice that *y*_0_ and *y*_1_ can be swapped to augment the training dataset. This self-supervised loss can be related to its supervised counterpart. For completeness, we give a standard derivation in Appendix B.

### 2.3 Missing wedge prediction

The data-fidelity loss enforces consistency only within the visible wedge, leaving the missing wedge unconstrained. We follow (Chen *et al*., 2021) and introduce an additional loss term based on rotation equivariance, which serves to guide the reconstruction in the missing wedge. The proposed loss function integrates two complementary components. First, it enforces equivariance of the combined neural network and masking operator, *f*_*ϕ*_(*A*(·)), under 3D rotations. Second, it ensures reconstruction consistency between the two measurements, *y*_1_ and *y*_2_. Together, these terms promote a consistent filling of the missing wedge.

Intuitively, the equivariant loss is minimized if the neural network performs well at filling missing wedge on any rotation of the input volume. This principle is formalized through the following loss function:

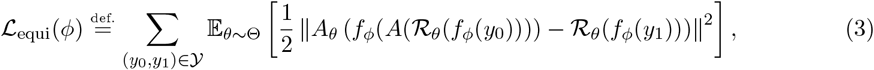

where ℝ_*θ*_ is the rotation operator by angle *θ* ∈ Θ, *A*_*θ*_ = *A* ◦ ℝ_−*θ*_ denotes the operator that masks the missing wedge rotated by *θ* and *θ* ∼ Θ is the uniform distribution over the finite set Θ. We take Θ to be the 20 fixed rotations that can be applied without interpolation, excluding rotations of 0° and 180° around the tilt axis, as well as their corresponding flip operations. In total, the set Θ contains 40 elements. The sampling procedure of rotations is similar to what is proposed in IsoNet, with the addition of flip operations to enhance data diversity. Similarly to the denoising loss, the role of *y*_0_ and *y*_1_ can be swapped to augment the training dataset. Ideally, the equivariant loss would be computed by replacing *f*_*ϕ*_(*y*_0_) and *f*_*ϕ*_(*y*_1_) by the clean volume *x*. Our closest estimate of the clean volume is the estimate produced by the neural network *f*_*ϕ*_ at its current weights.

The key difference between DeepDeWedge and Icecream is the double application of *f* in (3), which leads to a simpler, principled training protocol; a detailed discussion is in Section A.1.

### 2.4 Training

To reconstruct the volume densities, we minimize a total loss that combines the data-fidelity and equivariance terms,

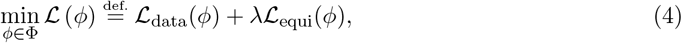

where *λ >* 0 is a regularization parameter to balance the importance of the two terms. We empirically found that *λ* = 2 provides optimal results, with strong denoising and without suppressing detail. An approximate solution of Problem (4) is computed using automatic differentiation and the Adam optimizer (Kingma, 2014).

### 2.5 Inference

The final reconstruction 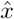 is computed by applying the trained neural network twice. First by computing the denoised tomograms and then by improving the missing wedge correction,

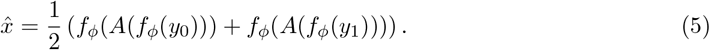

Applying the network only once leads to worse estimation of the missing wedge. This is because the equivariance loss, which is responsible for filling the missing wedge, is evaluated using the denoised volumes 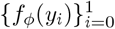.

To achieve optimal performance, the neural network needs to be trained on crops of a single tomogram and evaluated on the same crops. However, this requires training a separate network (for a few hours) for each tomogram. This process can be sped up if we are reconstructing multiple tomograms with the same biological content and acquired from the same microscope. In such cases, we can train a single network for all tomograms. As demonstrated in the next section, this network can even be effective on tomograms not used for training as long as they contain the same biological content and originate from the same microscope. When working with a structurally different tomogram, a network trained on multiple tomograms can be used as initialization for Problem (4) to speed up training; we show this in Section 3.4.1.

## 3 Results

We test Icecream on a variety of real datasets. We train the model with a patch size of 72^3^ voxels, for 50,000 iterations, with a batch size of 8, on a GeForce RTX 4090 GPU with 24GB memory using the Adam algorithm. Further details about the architecture and the training parameters used for the experiments can be found on the GitHub page of the project. We compare Icecream alongside with DeepDeWedge, Cryo-CARE+IsoNet, and CryoLithe. DeepDeWedge has been trained for 1000 epochs or 1 day, whatever limit is met first, with a patch size of 72^3^, using the code provided by the authors. IsoNet v0.3 has been trained for 30 epochs with a patch size of 72^3^ using the code provided by the authors. CryoLithe yields similar quality results than Cryo-CARE+IsoNet, but is significantly easier to apply as it does not require training a neural network. As a consequence, it can perform better than Cryo-CARE+IsoNet on tomograms that are difficult to process. For a tomogram of size 928 × 928 × 464, which is approximately the size of all the tomograms processed in this paper, Icecream is trained for 9 hours. On the same tomogram size, DeepDeWedge usually trains for 24 hours, and Cryo-CARE+IsoNet for 6 hours.

When the data can be split into four different tomograms, we train two independent models (Icecream, DeepDeWedge, or Cryo-CARE+IsoNet), each using two tomograms, and quantitatively asses the quality of the results using the Fourier shell correlation (FSC) between the two estimated tomograms. One major downside of self-supervised algorithms, and in particular of DeepDeWedge, is its long training time, on the order of dozens of hours. A direct implementation of Icecream suffers from the same drawback. To speed it up, we use mixed precision arithmetic on the GPU, enforcing the various quantities to be stored in half-precision (float 16). This significantly reduces the training time, without damaging the reconstruction performance. In the following, all Icecream results have been obtained using mixed precision.

### 3.1 Denoising and missing wedge correction of diverse cryo-ET datasets

We first showcase Icecream by reconstructing a variety of cryo-ET tomograms with different biological content and coming from different microscopes. The results are presented in Figure 1, with additional details in Table 1. Icecream performs equally well on different tomograms despite great structural differences. We observe stronger denoising and missing wedge correction on tomograms that have been obtained with well-aligned tilt-series; see Panel F in Figure 1. For comparison, Panel E shows a tomogram obtained from visually less accurately aligned tilt-series where reconstruction quality is not as good. The corresponding filtered backprojection (FBP) (Kak & Slaney, 2001; Harauz & van Heel, 1986) reconstruction can be found in the appendix in Figure C.1 and the corresponding DeepDeWedge reconstruction can be found in the appendix in Figure C.2. On the six tomograms, Icecream produces significantly sharper and better-denoised structures in X-Y plane compared to DeepDeWedge. In the X-Z and Y-Z planes, Icecream generally recovers membranes and particles with improved contrast and sharper definition, although in some cases the improvement is limited, as with the tomogram of panel E which suffers from imperfect preprocessing. A side by side comparison between Icecream and the reference methods is presented in the following paragraph.

**Table 1.** Information about the tilt-series used to generate Figure 1, Figure C.1 and Figure C.2.

| Image | EMPIAR ID | Tomogram | Resolution (Å) | Dimensions (px) | Split type |
| --- | --- | --- | --- | --- | --- |
| A | 11830 | 9 | 7.84 | $1024 \times 1024 \times 512$ | Dose |
| B | 11830 | 2 | 7.84 | $1024 \times 1024 \times 512$ | Dose |
| C | 12612 | 38 | 14.08 | $928 \times 928 \times 464$ | Dose |
| D | 11538 | 1435 | 4.0 | $1024 \times 1024 \times 512$ | Angle |
| E | 11658 | 1 | 7.84 | $1024 \times 1024 \times 512$ | Angle |
| F | 11058 | 3 | 14.08 | $928 \times 928 \times 464$ | Angle |

**Figure 1.**
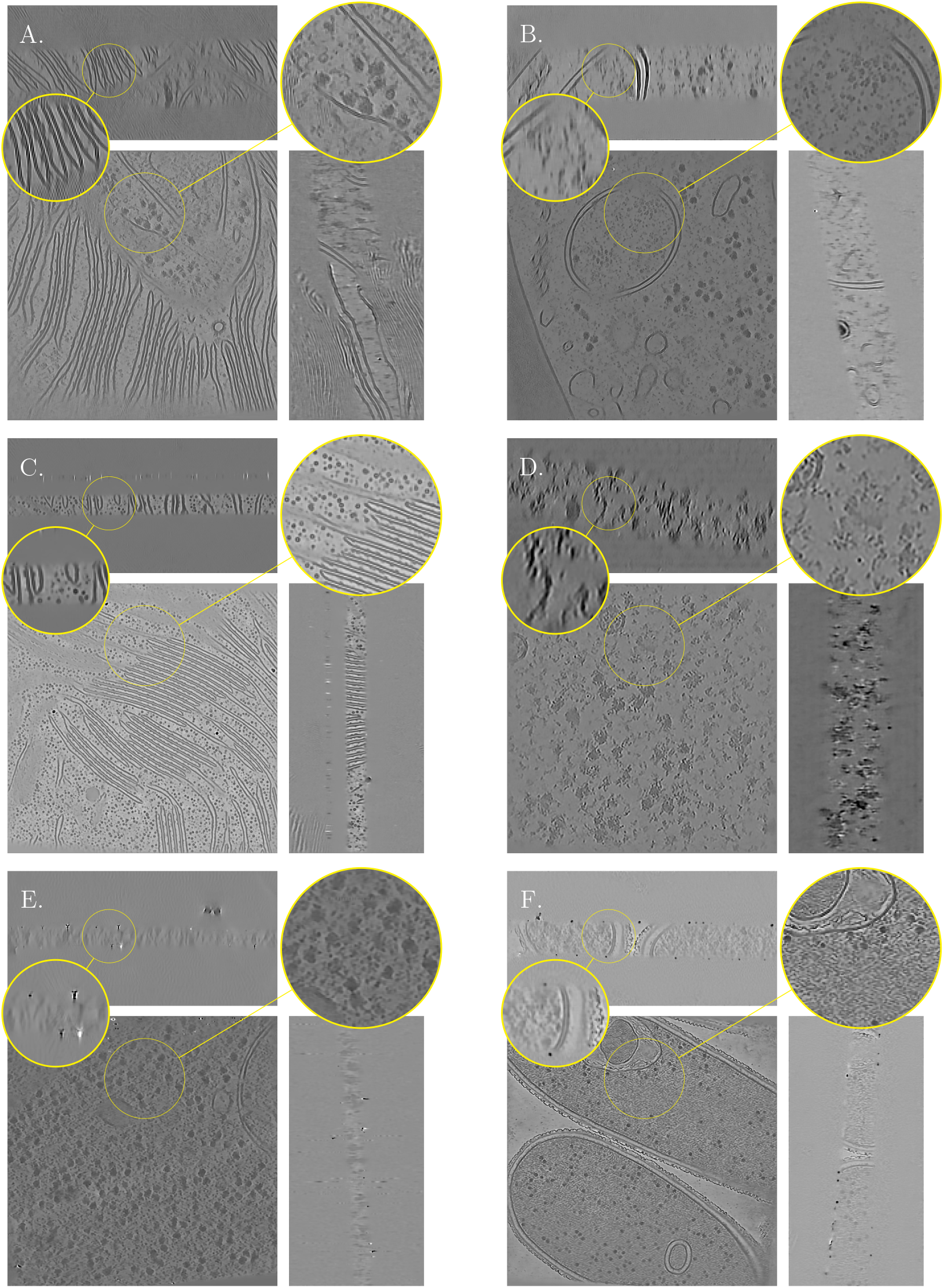
Icecream applied to a variety of tomograms, from different microscopes, different resolutions, and different tilt schemes. Information about the tomograms is reported in Table 1 Corresponding FBP and DeepDeWedge reconstructions are displayed, respectively, in Figure C.1 and in Figure C.2.

### 3.2 Post-processing of *T. kivui* tomograms

Next, we visually compare the proposed approach with the baseline methods on *T. kivui*, an anaerobic bacterium that efficiently fixates carbon (Dietrich *et al*., 2022). The raw dataset is available at the Electron Microscopy Public Image Archive (EMPIAR-11058). The post-processing results are displayed in Figure 2. We see that Icecream gives significantly better background denoising than the baselines, with empty regions appearing noticeably cleaner. It also makes it easier to discern small structures (e.g., orange arrow) and better recovers high-frequency patterns (e.g., red arrow). This indicates the potential of Icecream to facilitate manual or automated particle picking as a first step for sub-tomogram averaging.

**Figure 2.**
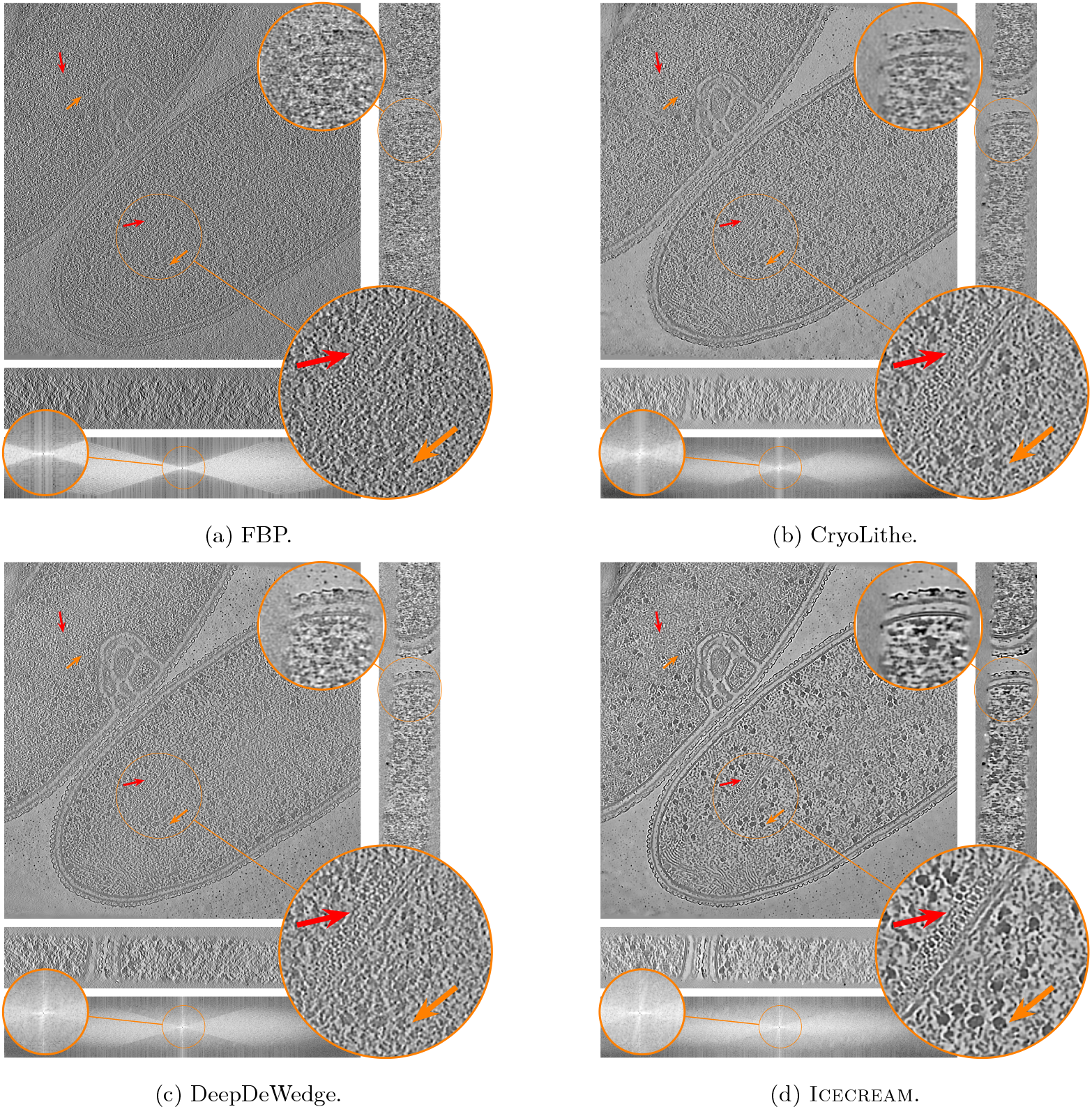
Orthogonal slices of the *T. kivui* (EMPIAR-11058) reconstruction using state-of-the-art denoising and missing wedge correction methods. Additionally, show the XZ slice of the Fourier transforms of the corresponding reconstructions in log-magnitude scale, where the observed and missing wedge supports are visible. Icecream performs better denoising, as visible in the background, and identifies sharper details, as indicated by the arrows. In the Fourier domain, Icecream shows stronger signal within the missing wedge region (brighter ares) compared to other methods.

### 3.3 Influence of splitting strategy

The proposed approach and the self-supervised baselines all require the input data to be split. The preferred way to do this is by splitting the dose to get two tilt-series. However, with older datasets or in some specific situations, this is not possible. An alternative solution is then to split a single tilt-series along the tilt angles, although this reduces the Fourier space coverage. In the following, we quantitatively evaluate the impact of splitting strategy on the different algorithms.

We work with the dataset of flagella of C. *reinhardtii*, which is the tutorial dataset used in cryo-CARE and contains 10 raw frames per tilt. The projections were collected at angles from −65^◦^ to +65^◦^ with 2^◦^ increments; pixel size is 2.36Å. The tilt-series is further downsampled by a factor of 6, resulting in an effective pixel resolution of 14.16 Å.

For the dose splitting evaluation, we use two frames per projection and average them at inference to construct the tilt-series. Frames 9 and 10 are excluded so that the four resulting tilt-series have comparable SNRs, assuming all frames contribute equally. We run DeepDeWedge, Cryo-CARE+IsoNet and Icecream independently on sets 1–2 and 3–4, producing two reconstructions. The two post-processed tomograms are then compared using FSC, with results reported in Figure 3a.

**Figure 3.**
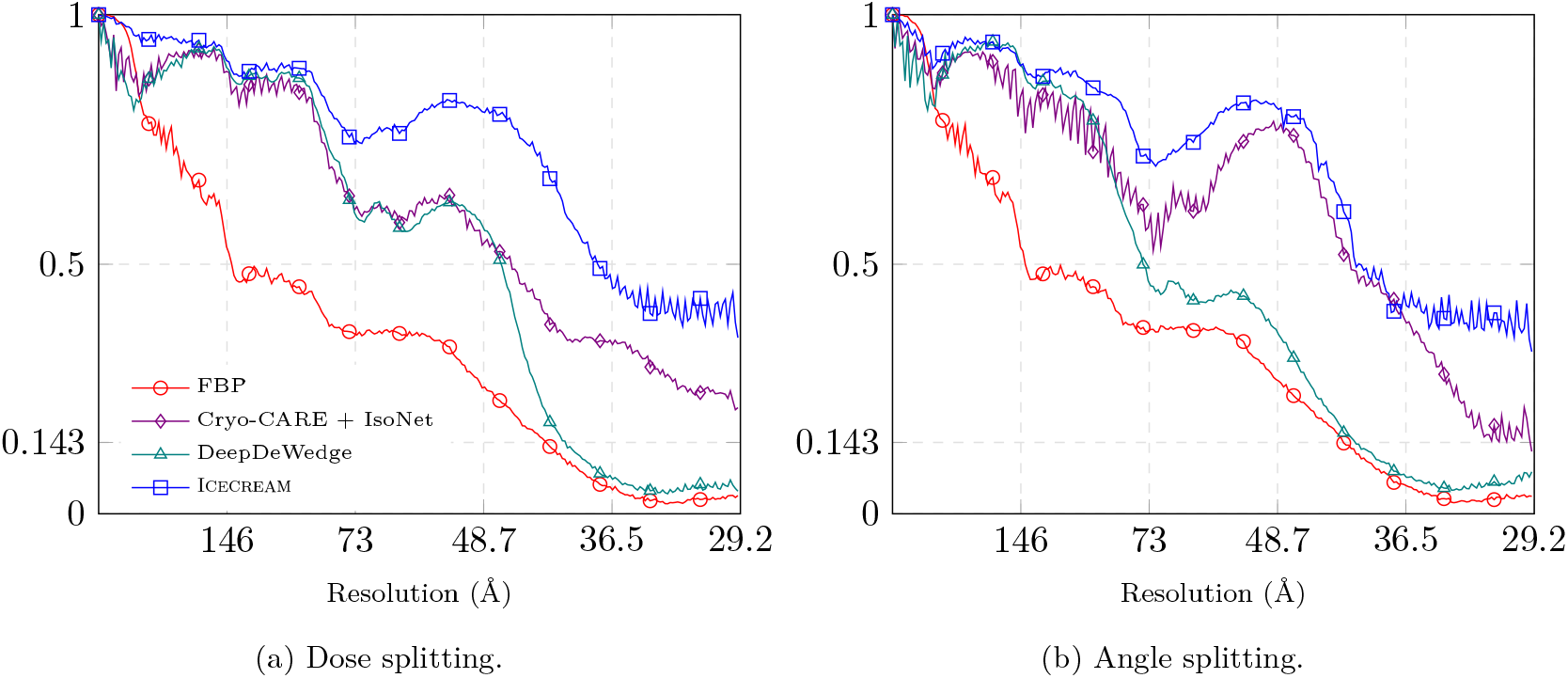
FSC of different methods on the dataset of flagella of C. *reinhardtii*. FSC was computed by splitting the data into 4 realizations, for which 2 pairs were used to produce two independent tomograms. Icecream performs significantly better than baselines uniformly on all frequencies and for both splitting strategies.

For angle splitting, we first average the 4 frames separately to obtain two tilt-series, again excluding frames 9 and 10 to have the same amount of information as for dose splitting. Each tilt-series is further split along tilt angles and fed into DeepDeWedge, Cryo-CARE+IsoNet, and Icecream. As before, we obtain two independent reconstructions and evaluate them using FSC; see Figure 3b.

Icecream outperforms the baselines methods for both splitting strategies. It is remarkable that the FSC for Icecream remains high even at high frequencies, though it should be interpreted with some care. It shows that the two statistically independent reconstructions obtained by Icecream are in better agreement than for other algorithms. This is in part a consequence of strong background denoising where other algorithms take a high-frequency FSC hit. Notwithstanding, an effective denoising method should remove noise in a consistent way across independent inputs, which is the case with Icecream. The corresponding processed tomograms are shown in Figure 4 (dose splitting) and in Figure 5 (tilt splitting).

**Figure 4.**
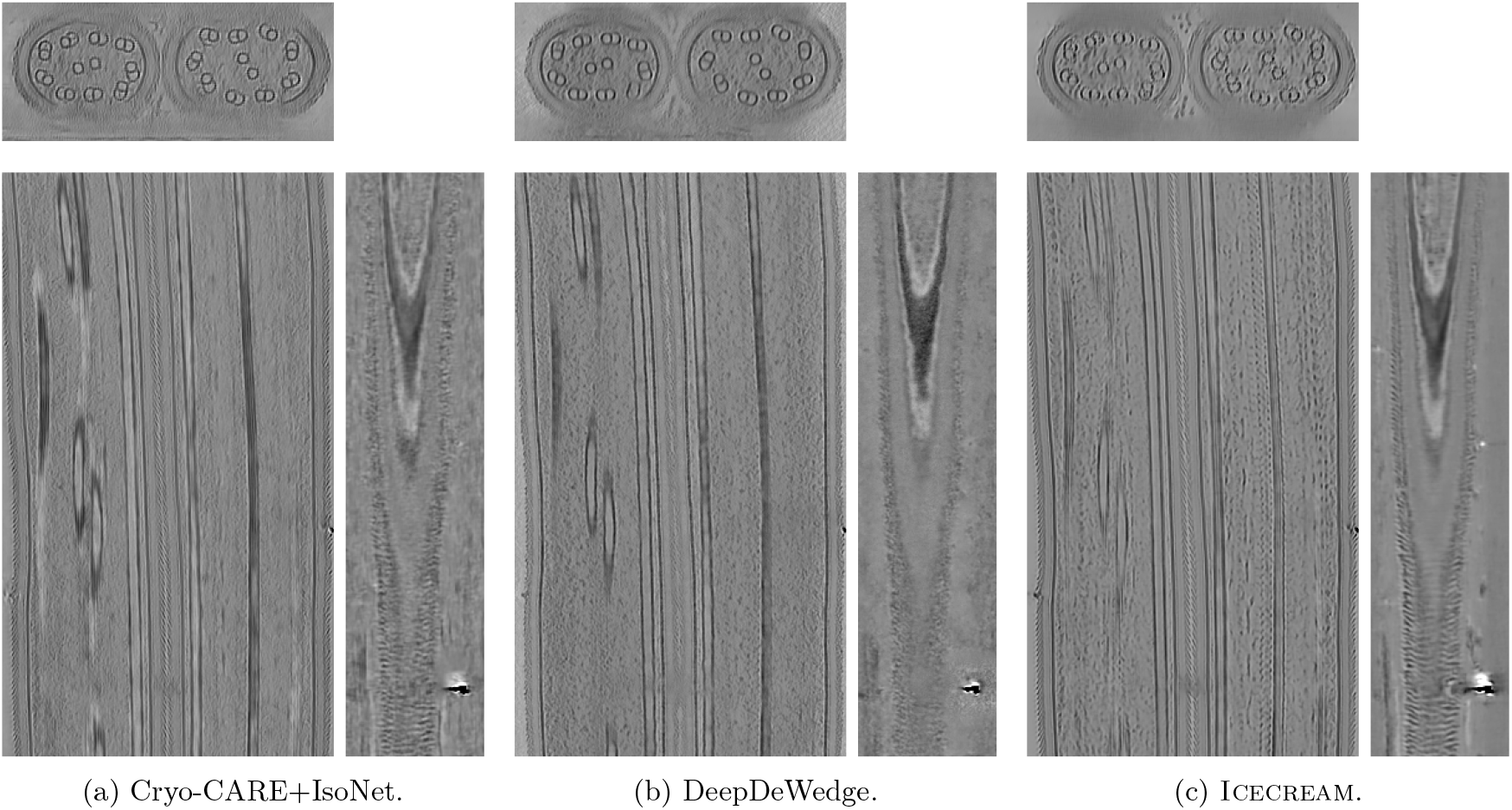
Orthogonal slices of flagella of C. *reinhardtii* recovered tomogram using *dose splitting*. The volumes have been cropped to remove empty areas.

**Figure 5.**
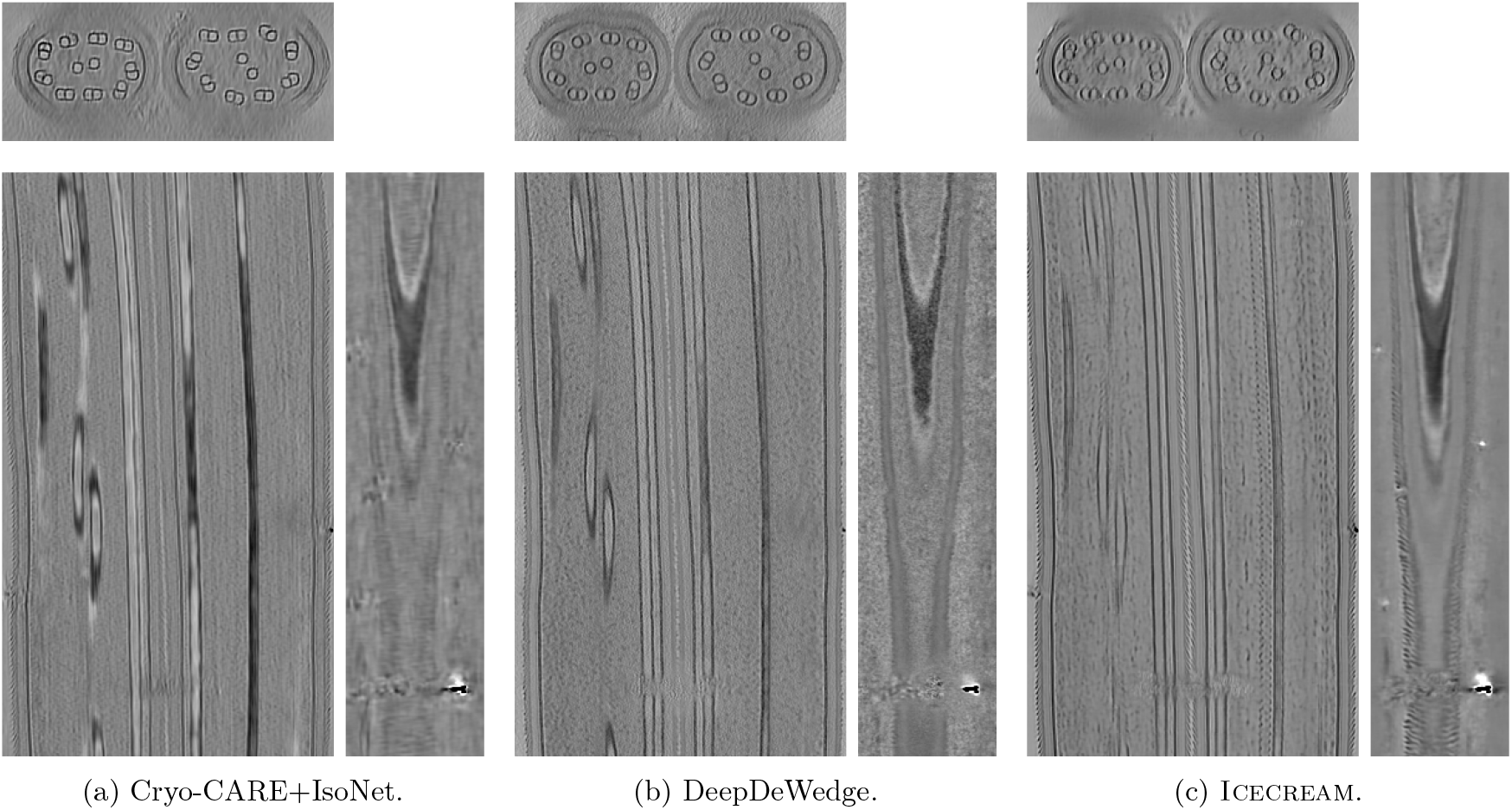
Orthogonal slices of flagella of C. *reinhardtii* recovered tomogram using *angle splitting*. The volumes have been cropped to remove empty areas.

Comparing Figure 3a and Figure 3b, it appears that dose splitting leads to slightly better performance. This result mirrors the conclusion of the Cryo-CARE paper that dose splitting should be preferred (Buchholz *et al*., 2019). In our case, the difference is relatively small; see Figure D.1 in Appendix for a side-by-side comparison for Icecream.

### 3.4 Performance of self-supervised methods on unseen data

One of the main limitations of existing self-supervised post-processing methods is that they require hours of training for each single tomogram. Here we explore using a network trained on a fixed dataset to process new tomograms. We show that such a network can be used without re-training if applied on data similar to the one it was originally trained on. If the tomogram to process is significantly different, a pre-trained network can still be used to speed-up the training procedure at the cost of finetuning.

We selected a subset of the EMPIAR-11830 dataset, which contains approximately 2000 tilt-series of *Chlamydomonas reinhardtii*. From this dataset, we chose 10 tilt-series for training and 4 for testing, all bin 4, resulting in a sampling rate of 7.84Å/pixel. The dataset provides dose-fractionated ODD and EVEN tilt-series. We used IMOD to generate the corresponding FBP reconstructions from the tilt-series that serve as inputs. During training, both ODD and EVEN volumes were used. However, at inference, we reconstructed the ODD and EVEN volumes separately on the test set. We evaluated the FSC between the ODD and EVEN reconstructions for Icecream and other baseline methods, see Figure 6. Since the volumes contain lamellae with varying thickness and tilt angles, we calculated the FSC on the central sub-tomogram of size 256 × 256 × 256.

**Figure 6.**
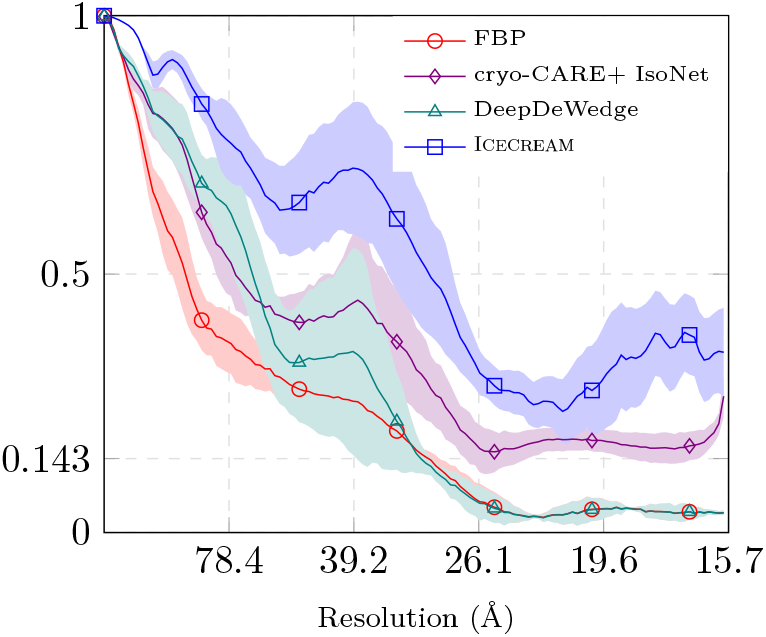
Out-of-distribution performance of self-supervised methods. The reconstructed volumes were obtained by using a neural network that has been trained on 10 different tomograms coming from the same microscope and containing similar biological content than the test tomograms. The FSC curve of Icecream consistently improves over FBP and improves over cryo-CARE+IsoNet and DeepDeWedge throughout the spectrum.

As with the in-distribution experiments, the FSC curve is consistently higher for Icecream than for the baselines. It is worse than in Figure 3, where the models were trained and evaluated on the same data, but the two experiments involve different datasets; on EMPIAR-11830 the FSC for FBP reconstruction is also worse.

The FSC gain obtained by using Cryo-CARE+IsoNet or DeepDeWedge instead of FBP is smaller for this experiment, which we again attribute to the fact that the models were not trained on this specific dataset. Icecream consistently improves over all baselines.

#### 3.4.1 Warm-start to accelerate processing

The performance of self-supervised models can degrade significantly when a pre-trained model is applied directly to a new dataset, especially if it is different from the training data. We now show that such a pre-trained model is nonetheless valuable as a warm start.

We use the neural network trained on the 10 volumes of the EMPIAR-11830 dataset as the pre-trained model. We demonstrate that fine-tuning this model for only 5,000 iterations yields reconstructions that are visually and quantitatively similar to those obtained by training Icecream from a randomly initialized network with 50,000 iterations and about 9 hours. Figure 8 shows the orthogonal slices of the reconstruction using only the pre-trained network, the network warm-started at the pre-trained weights, and the network started at random weights.

**Figure 7.**
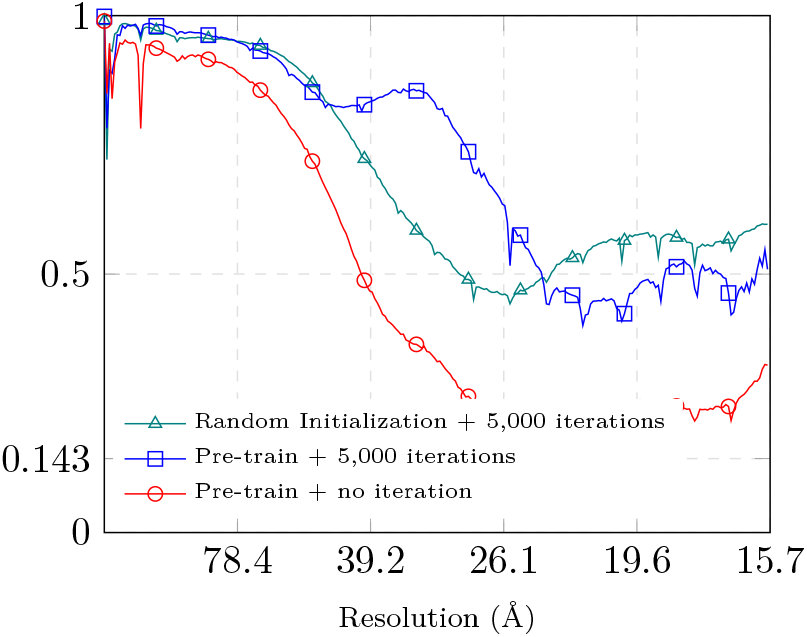
Influence of different finetuning strategies on the FSC. The reference volume is obtained by training the pre-trained model for 50,000 iterations. The FSC curves are computed using the final reconstructions of the respective models obtained after training for 50,000 iterations of as reference.

**Figure 8.**
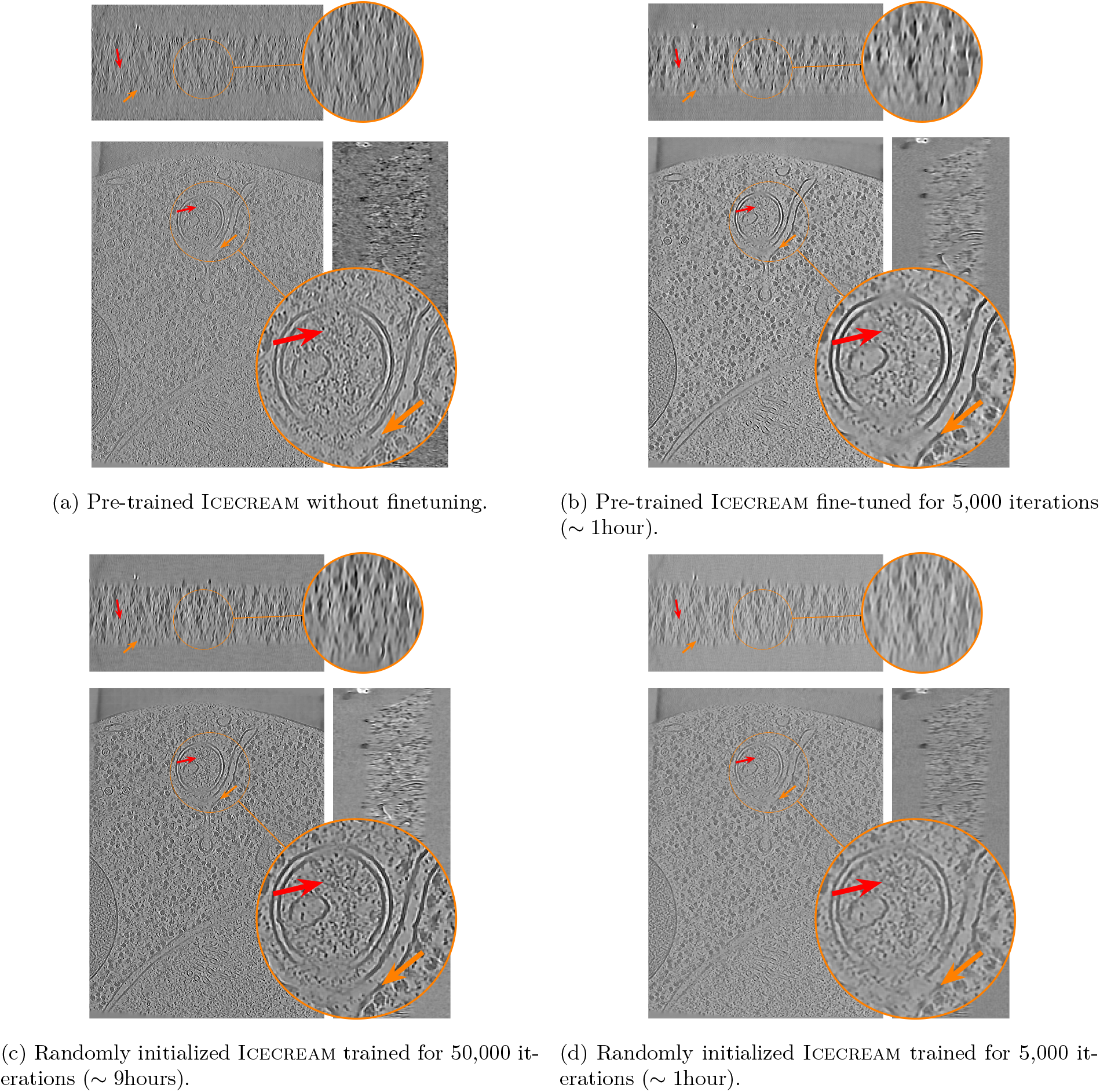
Inference of Icecream can be accelerated by fine-tuning a pre-trained network, reducing runtime from 9h to 1h. The models have been applied to a tomogram from EMPIAR 14162.

We observe that simply using the pre-trained network gives noisy reconstruction with missing detail. Remarkably, fine-tuning the pre-trained model for 5,000 iterations takes 1 hour and performs similarly to the randomly initialized model trained for 50,000 iterations and 9 hours. Finally, we observe that training a randomly initialized network for 5,000 iterations does not produce satisfactory results which means that pre-training is crucial. These observations are quantitatively validated by the FSC in Figure 7. The FSC curves are computed using the final reconstructions of the respective models obtained after training for 50,000 iterations as reference.

## 4 Discussion and conclusion

We showed that self-supervised learning grounded in equivariance attains state-of-the-art empirical performance in cryo-ET reconstruction; a number of theoretical and applied questions nonetheless remain.

Icecream assumes that the distribution of subtomograms is invariant under a subgroup of rotations. In practice this holds only approximately, so even with large datasets, expressive networks, and global optimization, the minimizer of the equivariant loss recovers an approximation to the estimator one would obtain with ground truth. It is therefore important to quantify (i) the deviations of real volumes from rotational invariance and (ii) how learning proceeds under this mismatch—especially because the equivariant loss alone admits trivial minimizers (arising from the double application of *f*_*ϕ*_). The importance of such analyses is amplified by the fact that deep models can hallucinate structure. Biologically salient findings should thus be corroborated with independent analyses, including the noisy but bullet-proof FBP. (We did not observe hallucinations in our testing of Icecream, but it seems naive to think that they are impossible.)

Training time remains a limitation because a model is currently trained per tomogram. Methods from the deep-learning literature, such as meta-learning (Tancik *et al*., 2021; Zhang *et al*., 2022), could help speed this up. We showed that a pretraining–fine-tuning strategy lowers reconstruction time by roughly an order of magnitude. Further exploration of cryo-ET foundation models—large pretrained backbones adapted with lightweight heads—seems to be a promising direction (Liu *et al*., 2024; Subramanian *et al*., 2023; Pyzer-Knapp *et al*., 2025)

An immediate possibility is to use Icecream reconstructions to curate training data for supervised methods like CryoLithe (Kishore *et al*., 2025)), which are much faster at inference. In practice one might switch between the two strategies depending on the type of analysis and observed performance.

We designed Icecream for practitioners who are not specialists in neural-network training: after installing a small set of dependencies, only minimal configuration is required to obtain high-quality reconstructions. The approach is not restricted to biological imaging. Electron tomography in materials science also suffers from the missing-wedge problem (often with higher SNR); in such cases, Icecream can be applied without modification.

## Acknowledgment

This project was supported by the European Research Council Starting under Grant 852821—SWING. V.D. is supported by the Agence Nationale de la Recherche (ANR) and the Ministère de l’Enseignement Supérieur et de la Recherche. Calculations were partially performed at sciCORE (https://sci-core.unibas.ch/) scientific computing center at University of Basel.

## A Differences between Icecream and existing self-supervised algorithms

The proposed approach is closely related to existing self-supervised methods for cryo-ET, namely DeepDeWedge and IsoNet. In the following, we describe these approaches and highlight the differences which lead to Icecream’s higher-quality reconstructions.

### A.1 DeepDeWedge

DeepDeWedge uses a loss function similar to the one used by Icecream, a variant of equivariance loss, but with several key differences. The two methods also use rather different pre-processing steps.

DeepDeWedge starts by pre-processing the full tomogram to correct for the CTF using IsoNet’s CTF deconvolution routine. It then extracts overlapping cubic sub-tomogram pairs, 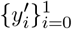, and normalizes them by batch to generate two sub-tomograms—the input of the model 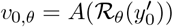 which is rotated and has a ‘manually’ removed missing wedge, and 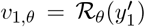, the rotated target.

#### Differences in the loss function

The DeepDeWedge loss is

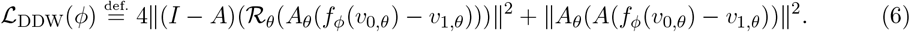

Differently from Icecream, DeepDeWedge uses a particular weighting applied in the Fourier space. The first term of the loss computes the equivariance loss with a stronger weight only on the span of (*I* − *A*)ℛ_*θ*_*A*_*θ*_ which is the (unrotated) missing wedge without the ‘manually’ introduced one in case they overlap. The equivariance loss is evaluated with a lower weight on the visible wedge (similarly corrected for the ‘manual’ one). Importantly, the equivariance loss in DeepDeWedge is different from the one used in Icecream in that it only applies *f*_*ϕ*_ once in the visible wedge. As a consequence, DeepDeWedge only uses one pass of the network at inference, while Icecream requires two passes, which we assume explains the improved quality of the denoising.

In Icecream, we evaluate this loss with the same weighting everywhere. The *θ*-rotated missing wedge is also removed, see Equation (3). This allows the regularization by equivariance to act everywhere in Fourier space. The importance of denoising in the visible wedge is controlled by the data-fidelity loss (2).

#### Iterative sub-volume updates

Similarly as in IsoNet, DeepDeWedge training uses a form of boosting: the entire training set of sub-volumes is periodically updated using the current trained network, and then treated as a new training set. As in IsoNet, this is done only on the missing wedge, while keeping the observed wedge untouched (and thus noisy).

Icecream is simply trained on the dataset by minimizing a principled loss, without boosting. This is motivated by the framework of equivariant imaging which suggests using the clean volume (Chen *et al*., 2021). Since the non-degraded volume is not available, we substitute the output of the network at the current iteration. The double application of the network in the equivariant loss plays a similar role as boosting in DeepDeWedge and IsoNet, but in a principled way which avoids the drawbacks of boosting. As explained in Section 2.5, it also means that we evaluate the network twice at inference. As explained in the previous paragraph, strong denoising is achieved in Icecream with the data-fidelity loss in Equation (2) which is not present in DeepDeWedge. The regularization enforced by the data-fidelity loss is fundamentally different than the one imposed by the equivariance loss (which rotates the sub-tomograms) and is similar to Cryo-CARE denoising (Buchholz *et al*., 2019).

#### Scaling-equivariant architecture

Similarly as in DeepDeWedge, we use an architecture inspired by the original U-Net. A key difference is that we design the network to be scaling equivariant: we use the 3D U-Net from Wolny et. al. (Wolny *et al*., 2020), but with biases removed from all convolutional layers and without normalization layers. This is very important in our setting: iterated application of the network to sub-tomograms changes spatial statistics between passes. See Appendix E for a related ablations. It has been showed by Mohan et. al. (Mohan *et al*., 2019) that scale equivariant networks are robust, interpretable and generalize well to unseen noise characteristics.

### A.2 IsoNet

IsoNet differs more substantially from Icecream and DeepDeWedge, but it shares the key characteristic that the dataset is augmented by rotating the sub-tomograms and removing the wedge. Similar to DeepDeWedge, the first step of IsoNet is to pre-process the input tomogram by compensating for the CTF. Unlike DeepDeWedge (but similarly to Icecream), sub-tomograms are extracted using a criterion that favors non-empty regions. At each iteration the sub-tomograms are rotated by randomly sampling among 20 rotations that require no interpolation; we adopt this strategy in Icecream.

An important advantage of IsoNet is that it does not require two observations of the same structure. This is circumvented by adding noise to the input of the network before removing the missing wedge. If *v* is the observed sub-tomogram, IsoNet’s loss function is

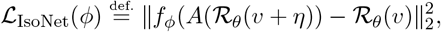

where *η* is Gaussian noise. The sub-tomogram *v* is iteratively updated by using the current neural network, *v* ← *f*_*ϕ*_(*v*). IsoNet’s loss is similar to the equivariance loss defined in Equation (3) modulo noise and the fact that it is evaluated on the entire Fourier space, including the original missing wedge. In practice, IsoNet is often combined with Cryo-CARE to boost denoising. In Icecream, the denoising step is directly included in the data-fidelity loss (2).

## B Self-supervised vs supervised denoising

In self-supervised learning we do not have access to ground truth training targets. It is nonetheless possible to establish certain equivalences between the supervised and self-supervised losses by exploiting statistical independence or symmetries of data and physics. For example, under our assumption that the measurements *y*_0_, 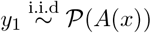 are independent and that

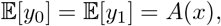

one can show that the minimizer of the expected data-fidelity loss is equal to the minimizer of its supervised counterpart,

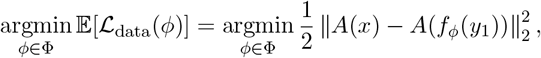

where the expectation is over *y*_0_. Indeed, following the standard derivation (e.g., (Lehtinen *et al*., 2018; Wiedemann & Heckel, 2024)), we have

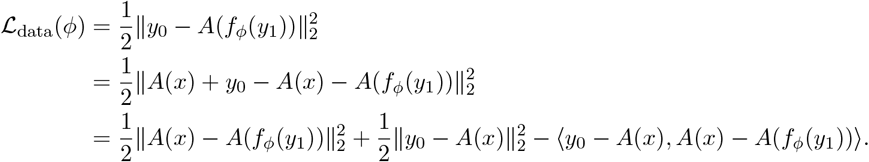

Taking expectation over *y*_0_ and using 𝔼[*y*_0_] = *A*(*x*), we get

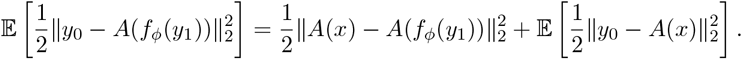

The only term depending on the optimization variable *ϕ* is 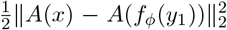, which is the supervised data-fidelity loss under the observation of *y*_1_.

## C Reconstruction of diverse cryo-ET datasets

We display the reconstruction of the volumes displayed in Figure 1 when using FBP reconstruction, see Figure C.1, and DeepDeWedge reconstruction, see Figure C.2.

**Figure C.1.**
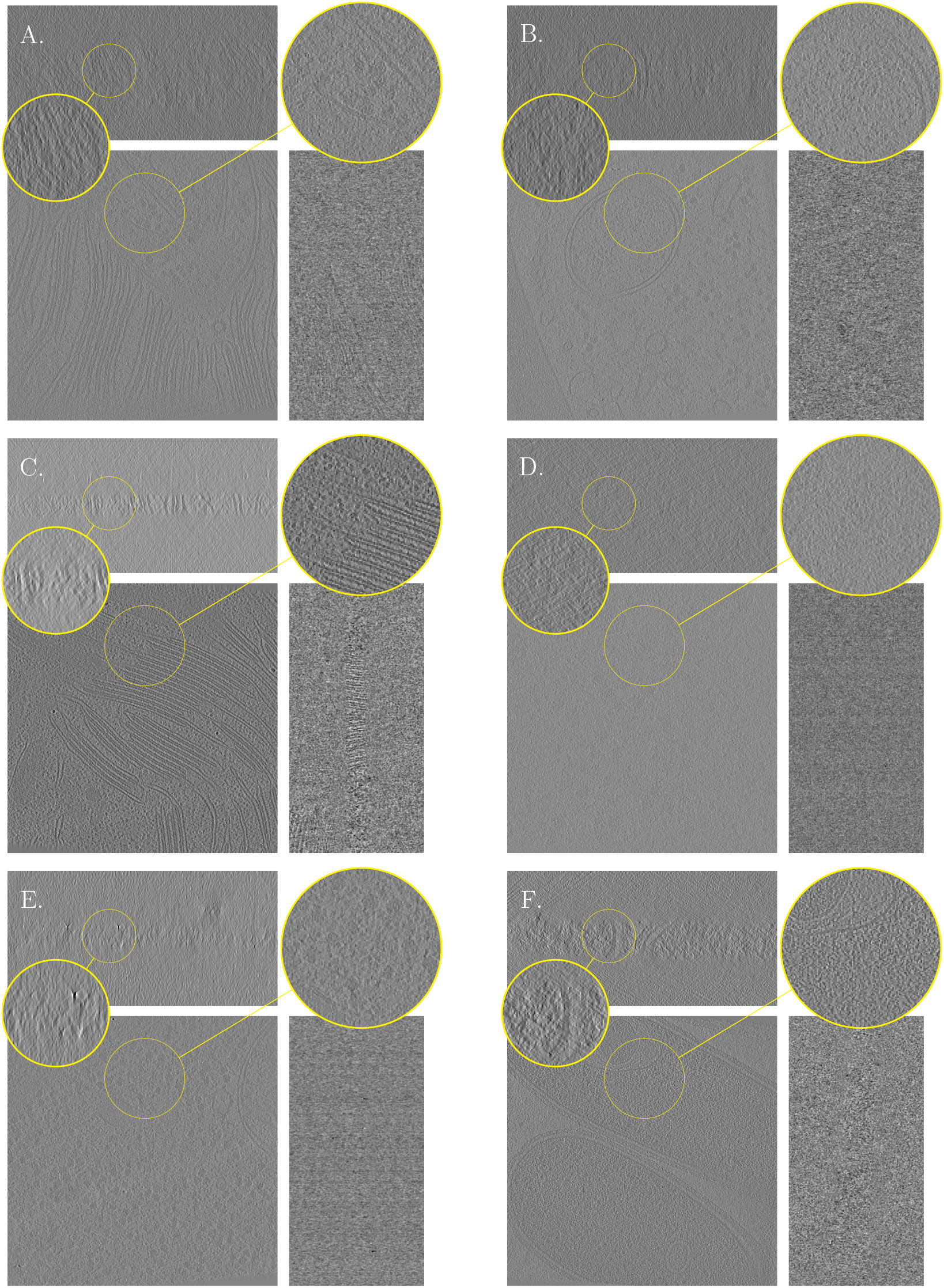
FBP reconstruction of the tomograms reconstructed in Figure 1.

**Figure C.2.**
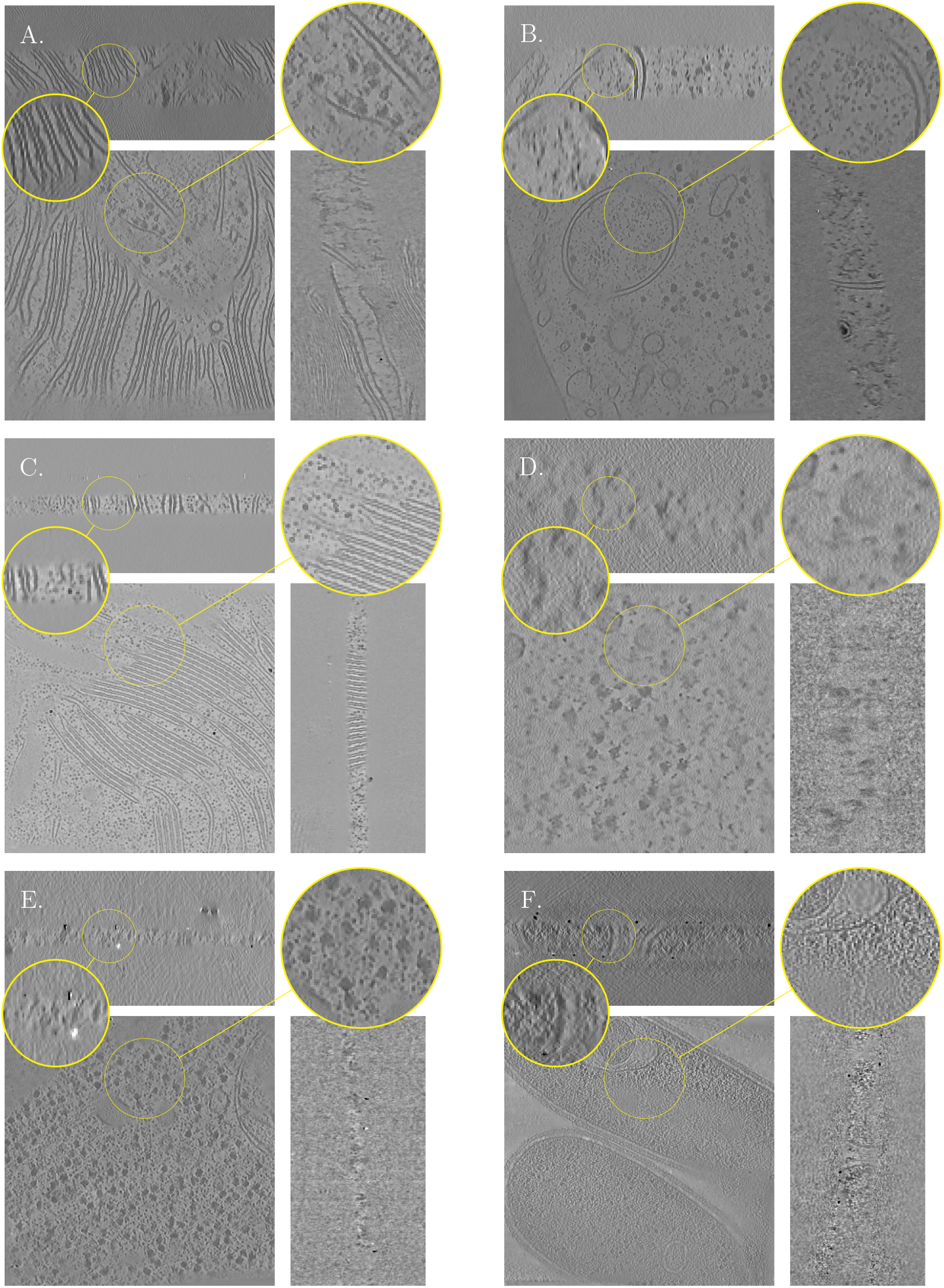
: DeepDeWedge reconstructions of the tomograms used in Figure 1.

For completeness with respect to Figure 2, we display the reconstruction of the *T. kivui* dataset using Cryo-CARE and Cryo-CARE+IsoNet in Figure C.3.

**Figure C.3.**
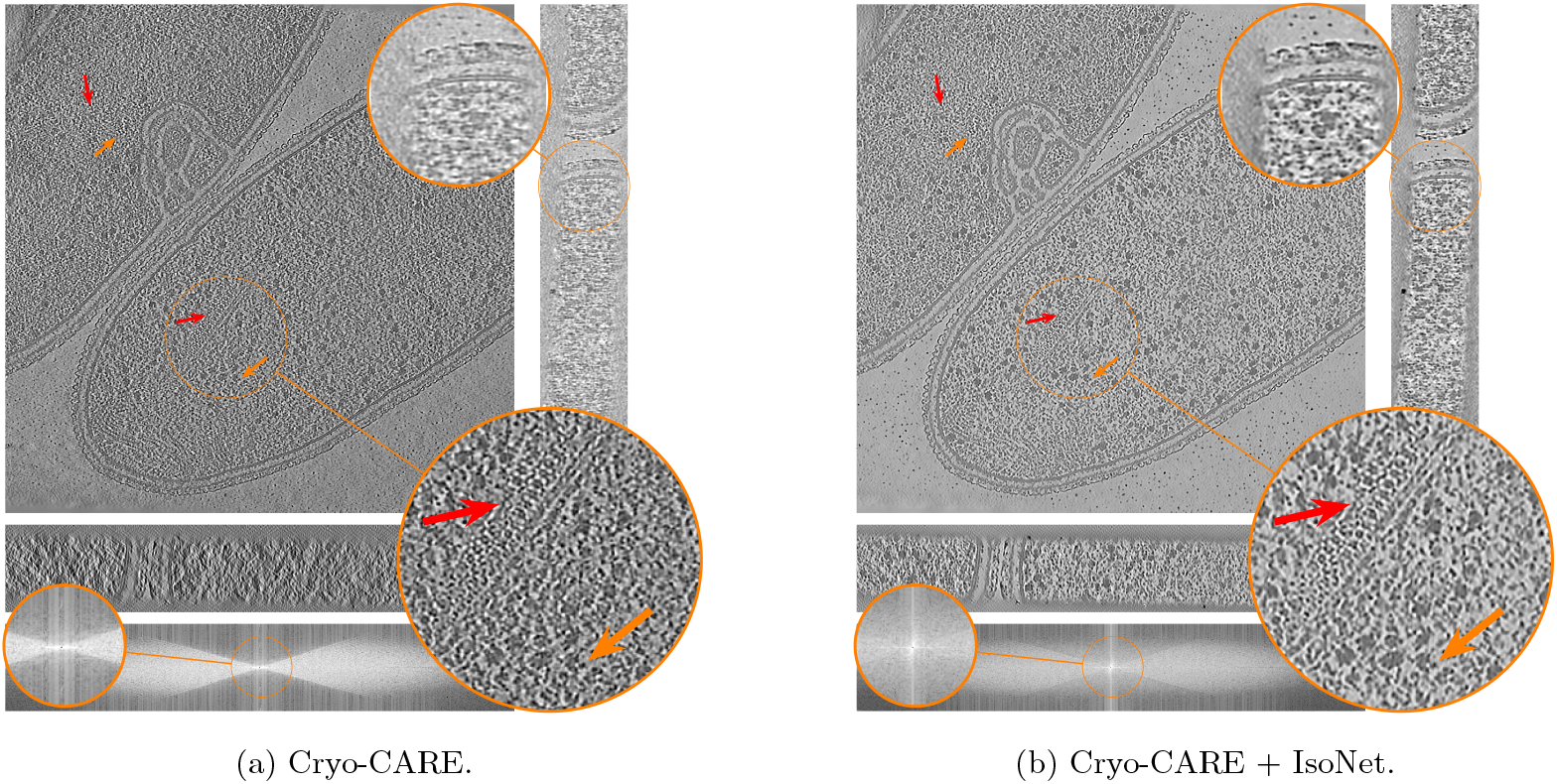
Orthogonal slices of the *T. kivui* (EMPIAR-11058) reconstruction using state-of-the art denoising and missing wedge correction methods. Baseline methods are displayed in Figure 2.

## D Dose vs tilt splitting

We compare the FSC curves of reconstructions obtained with Icecream when splitting the dose vs splitting the tilts, see Figure D.1. Even though the difference is small, dose splitting seems to produce better reconstructions in term of FSC.

**Figure D.1.**
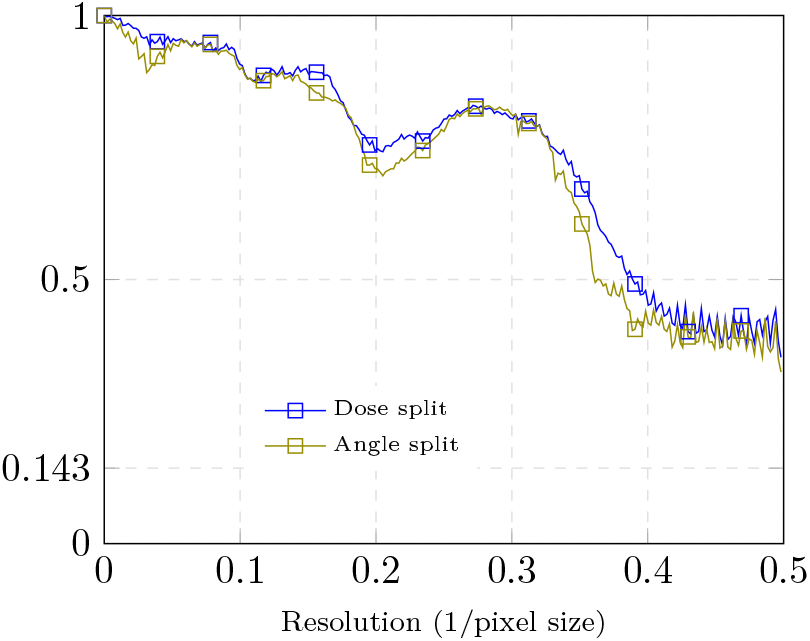
: Dose vs tilt splitting for Icecream on the dataset of flagella of C. *reinhardtii*.

## E Architectural Experiments

We experiment using the U-Net architecture from DeepDeWedge in Icecream method. Notice that the U-Net present in DeepDewedge is not scale equivariant, as opposed to the architecture of Icecream. We observed that removing learnable biases and instance normalization layers used in DeepDewedge’s U-Net implementation improves the reconstruction obtained when used in Icecream. We use the T. Kviu dataset where all the training parameters are kept constant except for the model architecture. Figure E.1 shows reconstructions using Icecream with various neural network architectures. We observed that removing normalization leads to the most noticeable improvement. Additionally, removing the biases further improves the quality of the reconstruction.

**Figure E.1.**
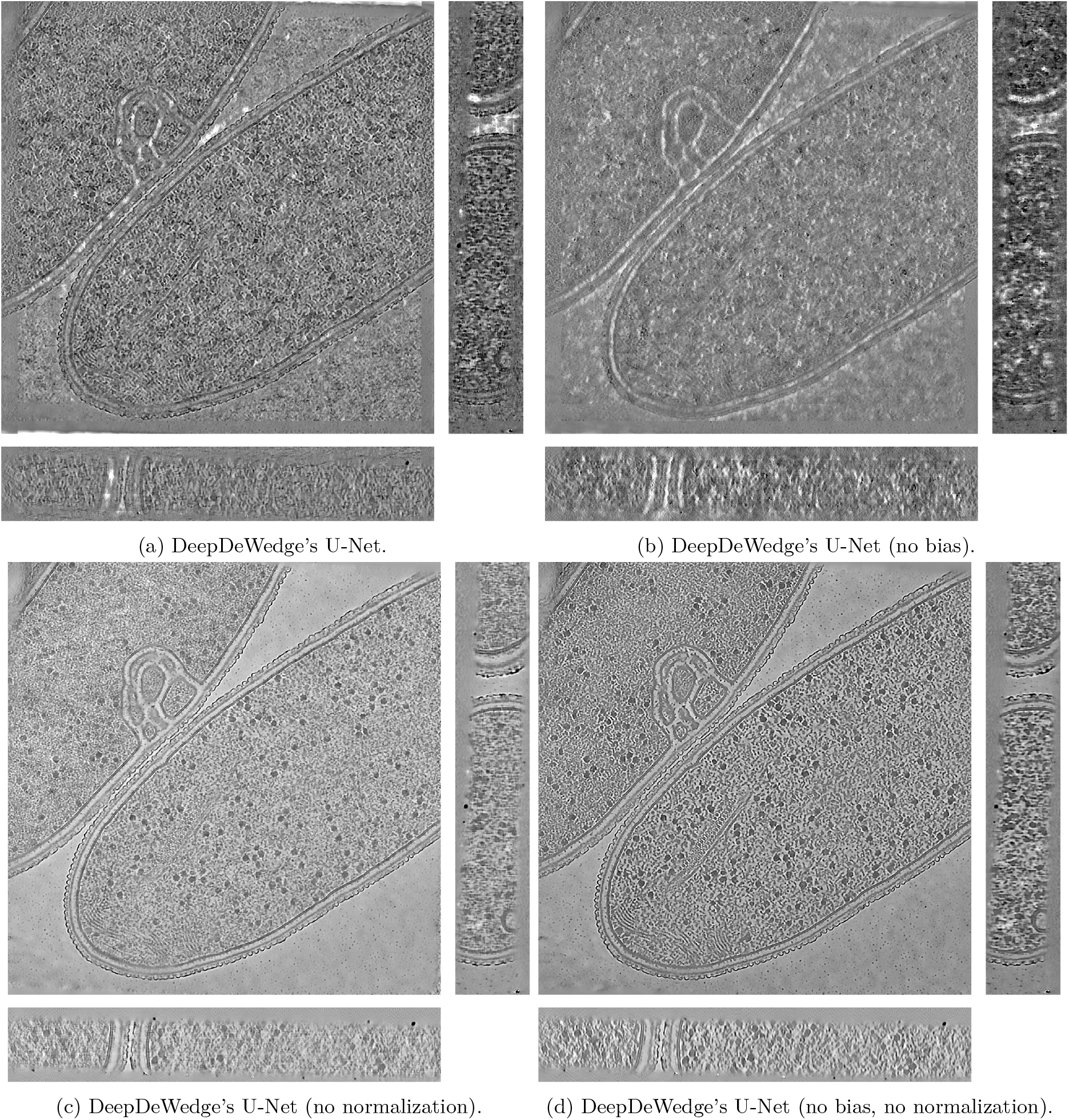
Reconstructions using Icecream with various neural network architectures. Results are obtained on the T. kivui dataset.

## Notes

https://github.com/swing-research/icecream

### Competing Interest Statement

The authors have declared no competing interest.

